# IL-1α induces cathepsin K in oral cancer cells – the invasion is unaffected

**DOI:** 10.1101/061465

**Authors:** Carolina C. Bitu, Ahmed Al-Samadi, Tuukka Alanärä, Hennaliina Karhumaa, Pirjo Viitasaari, Ivarne L.S. Tersariol, Fábio Dupart Nascimento, Tuula Salo

**Affiliations:** Cancer and Translational Research Unit, University of Oulu, Oulu, Finland.; Department of Oral and Maxillofacial Diseases, University of Helsinki, Helsinki, Finland.; Biochemistry/ Molecular Biology Department, Universidade Federal de São Paulo (UNIFESP), São Paulo, Brazil; Biomaterials Research Group and Biotechnology Division, UNIAN - SP, São Paulo, Brazil.; Medical Research Center Oulu, Oulu, Finland; Department of Oral Diagnosis, Oral Pathology Division, Piracicaba Dental School, University of Campinas, P.O. Box 52, 13414-903, Piracicaba, São Paulo, Brazil.; Helsinki University Hospital, HUSLAB, Helsinki, Finland

**Keywords:** Cathepsin K, OTSCC, IL-1α, CAF, HSC-3

## Abstract

**Background:** To study cathepsin K location in oral tongue squamous cell carcinoma (OTSCC), its traffic and expression levels in cultured oral cell lines; and to analyze the effect of interleukin (IL)-1α on OTSCC invasion.

**Methods:** Cathepsin K expression in OTSCC tissue samples was analyzed with immunostaining; its intracellular traffic was followed in HSC-3 cells after PMA treatment. HSC-3 and two oral keratinocyte cell lines, oral carcinoma associated fibroblasts (CAFs) and primary gingival fibroblasts (GF) were treated with IL-1α. Cathepsin K expression was measured using PCR and ELISA. Lastly, the effects of IL-1α on HSC-3 invasiveness, alone and in co-cultures with fibroblasts in the 3D myoma invasion model, were determined.

**Results:** Cathepsin K in OTSCC cells was found in vesicles close to cell membrane and within exosomes. While cathepsin K was expressed at the basal level in both epithelial cells and fibroblasts, basal IL-1α levels were higher in epithelial cells compared with GFs and CAFs. Cathepsin K expression was slightly induced by IL-1α in all cell lines, but it did not affect HSC-3 invasiveness.

**Conclusion:** In OTSCC, cathepsin K remains mostly intracellular and it is slightly secreted within exosomes. IL-1α treatment has no effect on HSC-3 invasiveness in 3D myoma model.

## INTRODUCTION

Oral cancer is the tenth most common cancer in men worldwide with over 40 000 new cases are expected to be diagnosed every year, in which over 50% of these cases are tongue cancers. Despite the advances in tongue cancer therapy including surgery, chemotherapy, and radiotherapy, the 5 years survival has not improve significantly with an average of 50% survival rate (1, 2).

Cathepsin K, a powerful collagenase, is one of the major players in the process of extracellular matrix turnover (3). Cathepsin K activity is also important for non-skeletal metabolism, both in normal and pathological conditions (4, 5, 6, 7). For instance, although in normal skin cathepsin K is mainly absent, it is found in dermal fibroblasts in recent surgical scars (3), being essential for the dynamic equilibrium between live mesenchymal cells and matrix synthesis and degradation (8).

Cathepsin K has also been studied in cancer as a major player in the ever-changing tumor microenvironment (TME), participating in the imbalance of extracellular matrix degradation, especially at the tumor invasive front. In skin squamous cell carcinoma (SSSC), cathepsin K in the stromal tissue facilitates cancer invasion (4, 5). Cathepsin K expression in fibroblasts was up-regulated through interleukin 1-alpha (IL-1α), which is secreted from SSCC cells in indirect co-culture experiments, showing that the cross talk between cancer and stromal cells are of paramount importance in understanding the complex process of tumor progression (5). In previous work, we reported the presence of cathepsin K in oral tongue squamous cell carcinomas (OTSCCs), with an association between expression pattern and prognosis (9). Down-regulation of cathespsin K significantly impaired OTSCC cell invasion *in vitro* 3D myoma model (9). However, the role of cathepsin K in the progress of oral carcinomas is still relatively unknown (10).

Since IL-1α was shown to up-regulate cathepsin K expression in SSCCs, this study aimed to investigate the role of IL-1α in OTSCCs, regarding mainly the regulation of cathepsin K expression, and its effects on cancer invasiveness. The location of cathepsin K was also studied within OTSCC tissue and cultured cancer cells.

## MATERIALS AND METHODS

### Ethics Statement

All patients signed an informed consent form and data inquiry was approved by the National Supervisory Authority for Welfare and Health (VALVIRA), #6865/05.01.00.06/2010 (5.10.2010), and the Ethics Committee of the Northern Ostrobothnia Hospital District, statement 49/2010 (16.8.2010). Usage of patient material for this study was approved by the Northern Ostrobothnia Hospital District Ethics Committee (statement #8/2006 and amendment 19/10/2006).

### Patient Samples and Immunohistochemistry

The 10 samples used in this study were selected to be most representative of the membranous expression pattern of cathepsin K, out of a previously described group of 121 OTSCC samples (9). Archival specimens of OTSCC samples, surgically treated at the Oulu University Hospital in 1981-2009, were retrieved from the Department of Pathology, Oulu University Hospital. Slides were immunostained for cathepsin K with the mouse anti-human cathepsin K IgG2b antibody (Biovendor, Candler, NC, USA) at a 1:500 dilution as described previously (9). For validation, we have used positive controls (human bone tissue) and negative controls (by omitting the primary antibody or using mouse primary antibody isotype control.

### Cell lines

HSC-3 tongue carcinoma cells (JCRB Cell Bank, National Institute of Health Sciences, Osaka, Japan), were cultured in 1:1 Dulbecco’s Modified Eagle Medium (DMEM) and Ham’s F12 culture medium supplemented with 10% fetal bovine serum. Gingival tissue-derived fibroblast cell line (GF) and carcinoma associated fibroblast cell line (CAF; ll) cells were cultured in l:l of DMEM with 10% fetal bovine serum. HMK (spontaneously immortalized oral keratinocyte cell line; 12) and IMHK (human keratinocyte cell line immortalized by transfection with the human papilloma virus-l6; l3) were cultured in keratinocyte serum-free medium (Gibco, Paisley, UK) supplemented with 0,005μg/ml recombinant human growth factor, 0,5mg/ml bovine pituitary extract and 100μM of CaCl2. THP-1 monocytic leukemia cells (American Type Culture Collection, Rockville, MD, USA) were cultured in RPMI culture medium supplemented with 25 μM HEPES, 100 units/ml penicillin, 1% L-glutamine, 50 μM β-mercaptoethanol and 10 % FBS.

### Immunocytochemistry and Phorbol ester-stimulated exocytosis

HSC-3 cells were cultured on round glass coverslips, placed in 6 well plates and incubated with 0,5µM LysoTracker^®^ (Life Technologies, Carlsbad, CA, USA) for 30 minutes in PBS. After, cells were incubated with 100ng/mL phorbol l2-myristate l3-acetate (PMA; Sigma-Aldrich, St. Louis, MO, USA) for different time points: 30 minutes, 1 hour, 2 hours and 3 hours. The immunofluorescence staining for cathepsin K was performed as follows: samples were washed five times with PBS and fixed in 2% paraformaldehyde in PBS for 30 minutes. After two PBS washes, samples were washed once with 0,1M glycine in PBS and twice in PBS. Cell permeabilization was performed in 0.0l% Saponin l% BSA solution in PBS for 10 minutes at room temperature. The primary antibody incubation was done in 1:250 in Saponin/BSA solution for one hour. Samples were given sequential washes with PBS solutions of 0.01% Saponin, 0.01% Saponin/1% BSA and again with 0.01% Saponin. After the secondary Alexa Fluor^®^488-conjugated antibody incubation at 1:500 in PBS with 0.01% Saponin, samples were washed four times with PBS. Samples were then counterstained with 4’,6-Diamidino-2-Phenylindole, Dihydrochloride (DAPI) for 15 minutes and mounted in glass slides. The labeled molecules in acidic compartments were analyzed using inverted confocal laser-scanning microscopy (Zeiss LSM-780, Carl Zeiss, Jena, Gernany)

### Human Myoma Organotypic Culture, Fluorescent Labeling and Quantification of Invasion Results

The myoma organotypic experiment was done as described previously (14), with few modifications. In preparation for the experiments, 9 myoma disks were incubated overnight in full DMEM-F12 1:1 culture medium and other 9 myoma disks were incubated in the same medium with the addition of 5ng/ml human recombinant IL-1α (Sigma-Aldrich). The myoma disks were then placed into Transwell^®^ inserts (Corning, NY, USA). A few minutes before plating, HSC-3 cells were labeled, according to manufacturer’s instructions, with the red lipophilic carbocyanine dye, Dil Vybrant™ (Life Technologies). GF and CAF cells were labeled with the green dye DiO Vybrant™ (Life Technologies). Labeled HSC-3 cells were plated on top of the myoma disks alone or in co-culture with either GF cells or CAF cells, in a total amount of 4×10^4^. All experiments were performed in triplicate. Cells were allowed to attach overnight and moved to uncoated nylon disks on curved steel grids in 12-well plates. One ml of medium with or without IL-1α was added to each seeded myoma disk. During the experiment, fresh media was added every third day, until the tenth day. Then the disks were fixed in 4% paraformaldehyde for 20 minutes at room temperature. Subsequently, for best preservation of morphological features, samples were immersed in a sterile 30% sucrose/PBS solution overnight at 4°C. The myoma disks were embedded in TissueTek^®^ OCT compound (Sakura, Alphen aan den Rijn, The Netherlands) and snap frozen in liquid nitrogen and kept at −70°C until use. Serial sections of 10μm were either stained with monoclonal mouse antihuman pancytokeratin IgG (clones AE1/AE3) at a 1:250 dilution (Dako, Copenhagen, Denmark) or counterstained with DAPI and kept protected from light until their use in the multicolor fluorescent imaging.

The fluorescent samples were then digitally recorded with the Zeiss LSM 780 Laser Scanning Confocal Microscope (Carl Zeiss Microscopy, Oberkochen, Germany). Immunostained samples were digitally photographed at x100 magnification (Leica Microsystems, Heerbrugg, Switzerland). Invasion depth was measured as the distance between the surface of the myoma tissue and the deepest invading cell in each sample. The invasion index was calculated as 1-[noninvading area/total area] using the measured areas of invading cells and noninvaded myoma tissue of each sample, excluding nonuniform invasion at the borders of the myoma, as described previously (14, 15).

### IL-1α experiment

Cells were cultured until 80% confluence, 1x106 cells were plated for each experiment. Cells were allowed to attach for 24 hours and starved of serum or cell growth supplements for additional 24 hours. After that, cells were treated with 5ng/ml recombinant human IL-1α in opti-MEM medium (R&D Systems, Minneapolis, MN, USA) for additional 24 hours.

### Total RNA extraction

Cells were washed once with chilled PBS and 1.5 ml of the commercial mixture of guanidine thiocyanate and phenol, TRI reagent^®^ (Sigma-Aldrich) was added. RNA extraction was performed according to manufacturer’s instructions. Samples were resuspended in RNase free water up to 30 μl. Total RNA quantitation and purity were determined through spectrophotometry in NanoDrop instrument (Thermo Scientific).

### Reverse transcriptase polymerase chain reaction (RT-PCR)

Complementary DNA (cDNA) was synthesized using 1 μg of total RNA per sample. The reaction was performed using 200 U of the reverse transcriptase RevertAidTM (Fermentas-Thermo Scientific, Fremont, CA, USA), 20 U of the RNase inhibitor RiboLockTM, 1mM of dNTP mix (Fermentas-Thermo Scientific) and 100 pmol of random hexamers (Fermentas-Thermo Scientific). Due to the use of random hexamers, the reaction was designed to include an incubation period of 10 minutes at 25 °C followed by 60 min at 42 °C for cDNA synthesis and 10 minutes at 70 °C for enzyme inactivation.

### Polymerase chain reaction (PCR)

The PCR reactions were performed with primer sequences specific for the amplification of the following genes: cathepsin K primers (forward 5’-ccgcagtaatgacacccttt-3’ and reverse 5’-gcacccacagagctaaaagc-3’), IL-1α (forward 5’ - aatgacgccctcaatcaaag-3’ and reverse 5’ - tgggtatctcaggcatctcc-3’), TGF-β1 (forward 5’-gtggaaacccacaacgaaat-3’ and 5’-reverse cacgtgctgctccactttta-3’) and beta-actin (forward 5’-aactgggacgacatggagaaaa-3’ and reverse 5’-ag aggcgtacagggatagcaca-3’) which was used as a reference gene. Briefly, PCR reactions were done in 45 cycles of 5 minutes at 95 °C, 2 minutes at 94 °C, 1 minute at 54 °C, 30 seconds at 72 °C and a final extension at 72 °C for 10 minutes, using the DNA polymerase AmpliTaqGold^TM^ (Fermentas). The PCR products were then separated by electrophoresis in a 1% agarose gel.

### Total protein extracts

Cells were scraped from cell culture plates with the addition of ice cold lysis buffer (50 mM Tris, l0 mM CaCl2, l50 mM NaCl, 0.05% Brij, pH 7.5). The collected cells in the lysis buffer were kept under rotating agitation overnight at 4 °C. Following this incubation, the supernatant was separated from the insoluble precipitate by centrifugation at 12000 x g for 10 min at 4 °C. Protein extracts were then kept in −70 °C until use. Protein concentrations were measured through the Lowry colorimetric assay, Bio-Rad DC protein assay (Bio-Rad, Hercules, CA, USA).

### Exosome isolation

HSC-3 cells were washed with PBS, and cultured in Opti-MEM^®^ I Reduced Serum Medium (Life Technologies) with the addition of 50 μM 4-aminophenylmercuric acetate (APMA; Sigma-Aldrich), as described previously (16). Incubation lasted for 6 h, after which the media were collected and the exosomes were isolated with ExoQuick-TC^TM^ kit (Systems Biosciences, CA, USA) according to the manufacturer’s instructions. Briefly, the ExoQuick-TC solution was added to the supernatant in a 5:1 ratio. This sample/ExoQuick solution was incubated at 4°C overnight. The solution was then centrifuged 1500 × g for 30 min and the supernatant was removed. The precipitate was suspended in 1 × RIPA-buffer (25mM Tris-HCl pH 7.6, 150mM NaCl, 1% NP-40, 1% sodium deoxycholate, and 0.1% SDS) for 5 minutes in room temperature for Western blotting.

### Western blot

Western blot analysis using a murine monoclonal cathepsin K antibody in a 1:1000 dilution (Biovendor) was performed as described previously (15). As a positive control for cathepsin K western blot, total protein extract from THP-1 cells was used.

### Enzyme-linked quantitative immunoassay (ELISA) for cathepsin K

Cell homogenates and media samples were measured for their cathepsin K content using the ELISA kit composed of polyclonal sheep anti cathepsin K antibody coated microtiterstrips (Biomedica Gruppe, Vienna, Austria). Fifty μl of undiluted sample was added to each well. Standard curves were calculated according to manufacturer’s instructions. Cathepsin K final quantitation was calculated as a ratio relative to total protein content from each sample.

### Statistical analysis

Kruskall-Wallis test was used to check the statistical significance of grouped data. Values of *P* < 0.01 were considered as statistically significant.

## RESULTS

### Cathepsin K is not located solely in acidic lysosomal subcellular compartments in OTSCC cells

*In vivo*, some OTSCC cells presented a pattern of cathepsin K accumulation in cytoplasm, whereas in other cases it was accumulated in proximity of the cytoplasmic membrane. We hypothesized that, although we could not see positive immunohistochemistry staining for cathepsin K in the extracellular matrix, it still might be secreted from OTSCC cells into the TME within exosomes (9; Figure 1A). Therefore, we performed an experiment where we used PMA, a known tumor promoter, to induce cell stress and lysosome purging. In our experiments, at the initial time point, HSC-3 cells showed that the majority of cathepsin K (stained green) co-localized with lysosomes (labeled red; Figure 1B). In other time points of PMA treatment, there was a noticeable scattering of cathepsin K throughout the cytoplasm. The same is not true for the lysosomes, which also migrated, although to different directions, as clearly demonstrated (last three panels of Figure 1B). These results demonstrate that cathepsin K vesicles are carried towards the cellular membrane, similarly to what is shown in the *in vivo* samples (Figure 1A). However, by these methods we could not identify cathepsin K in the extracellular spaces. Instead, in a complementary investigation, we found that in cultured HSC-3 cells, small amounts of cathepsin K were actualy found in exosomes secreted into culture media (Figure 2A).

**Figure 1.**
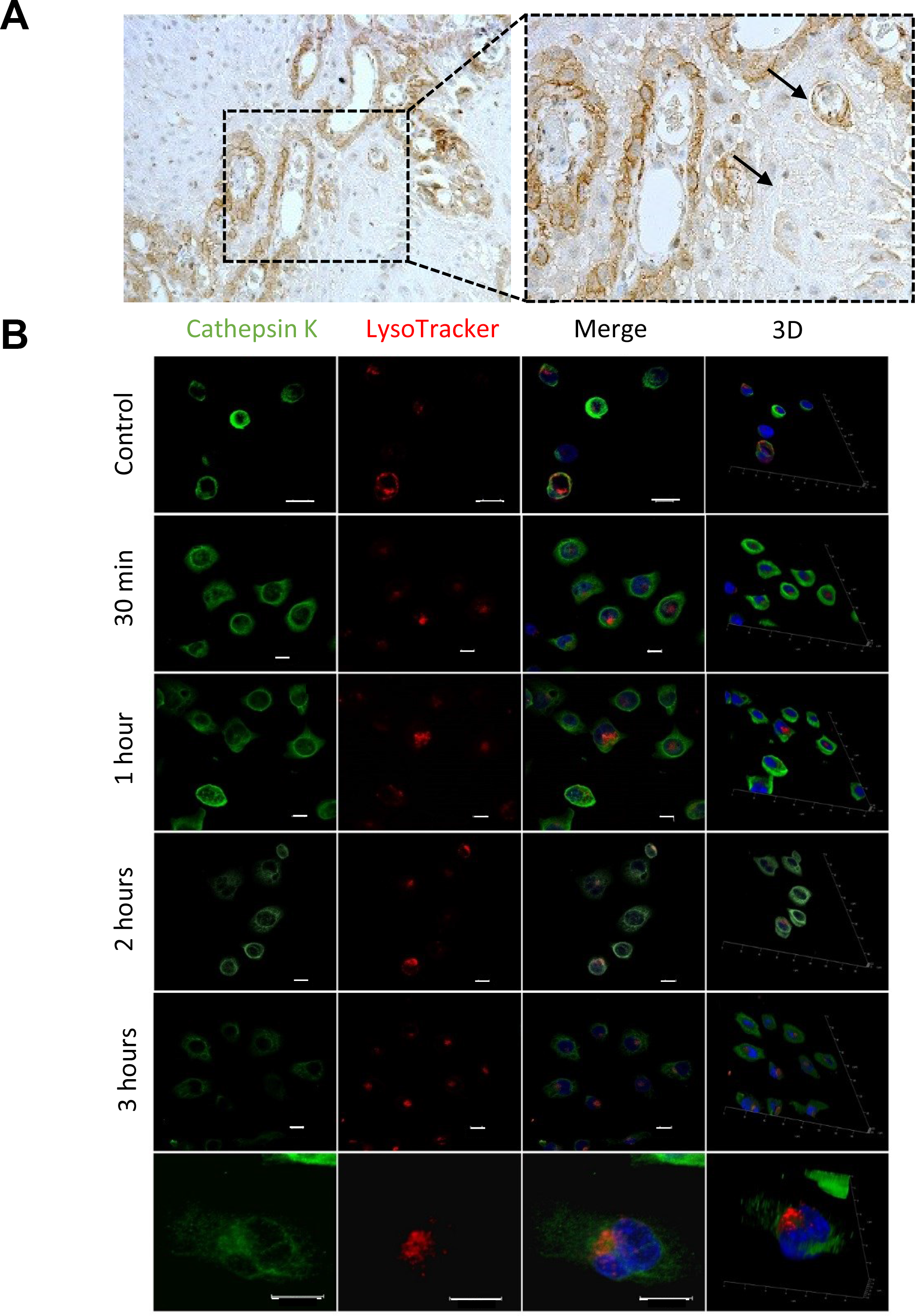
*In vivo* analysis of cathepsin K localization and intracellular traffic. **(A)** In immunohistochemistry analyses *in vivo*, some OTSCC cells present a very distinct localization of cathepsin K droplets close to the cytoplasmic membrane, suggesting that cathepsin K may be secreted (arrows). **(B)** In the confocal immunofluorescence imaging of OTSCC cells *in vitro*, there is co-localization of lysosomes and cathepsin K in control cells. However, when cells were treated with PMA, cathepsin K (green) travels towards the membrane, indicating that cathepsin K is being transported but not secreted by OTSCC cells. Last horizontal panel shows an individual cell after 2 hours of treatment with PMA. Cathepsin K vesicles begin to travel towards the cell membrane, while lysosomes gather close to the nucleus. Scale bars =10 μm.

**Figure 2.**
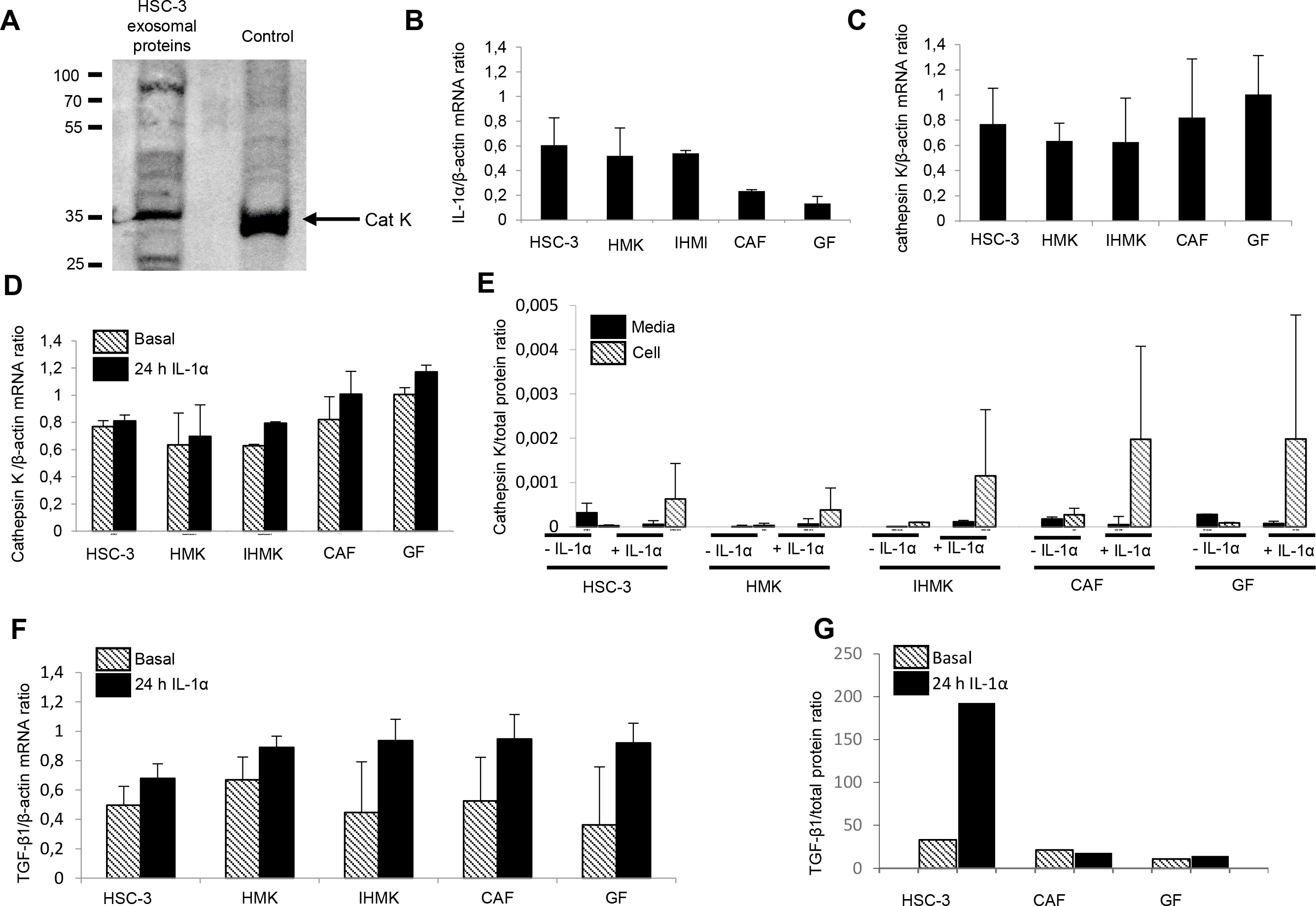
Cathepsin K levels are variable between epithelial and mesenchymal cell lines and presence of exogenous IL-1α. **(A)** Extracellular cathepsin K was found in proteins isolated from purified exosomes extracts from HSC-3 cell media. Total protein extract from THP-1 cells is used as positive control. **(B)** Basal IL-1α levels were higher in HSC-3, IHMK and HMK cell lines than in fibroblasts (CAF and GF). The results are shown as relative to expression of the reference gene, β-actin. **(C)** Cathepsin K basal mRNA levels vary, with no significant difference between the cells lines. The HSC-3 line, CAF and GF present slightly higher levels of expression than HMK and IHMK cell lines. The reference used for gene expression levels normalization is β-actin. **(D)** Exogenous IL-1α was able to induce expression of cathepsin mRNA in all cell lines, compared to basal levels **(E)** After induction of cathepsin K expression by IL-1α, the amount of cathepsin K both in cell extracts or secreted in media was measured. In cell extracts, the levels of cathepsin K were very low, however, after IL-1α treatment, there was a remarkable increase, especially in the CAF and GF cell lines. In media, cathepsin K levels were initially low in HSC-3, CAF and GF and virtually inexistent in the HMK and IHMK cell lines. After IL-1α induction, levels even dropped slightly for the HSC-3, CAF and GF cell lines. Total cathepsin K levels are shown as relative to total protein content.

### Cathepsin K basal mRNA levels are variable between all cell lines

The mRNA levels for cathepsin K in all of the cell lines did not differ significantly neither when compared individually nor when analyzed in groups of epithelial or mesenchymal origin. However, these results demonstrated that, at basal levels, cathepsin K mRNA levels were slightly higher in HSC-3, CAFs and GFs, than in HMK and IHMK (Figure 2C).

### Cathepsin K increased expression is mediated by IL-1α

The original hypothesis of this investigation proposed that IL-1α would mediate cathepsin K regulation in the cross-talk between oral epithelial and mesenchymal cell lines, like in the case of SSCC (17). First, we assessed IL-1α mRNA for each cell line. Our results showed that mesenchymal cells (CAFs and GFs) expressed low levels of IL-1α mRNA. Oral epithelial cell lines (HMK and IHMK) and the OTSCC cell line HSC-3 expressed more IL-1α mRNA (Figure 2B). We, then, cultivated all cells with media supplemented with IL-1α for 24 hours, and found out that, in all cell lines, there was a slight increase, in cathepsin K mRNA levels (Figure 2D).

### Cathepsin Kprotein expression levels are elevated by exogenous IL-1α

After supplementing culture media with IL-1α, we also assessed cathepsin K protein levels with ELISA, from cell lysates and culture media samples. Initial cathepsin K levels in cell lysates were quite low (Figure 2E) and the highest level was found in the CAFs. After IL-1α treatment, there was a noticeable increase in the amount of cathepsin K levels. The highest average cathepsin K level was observed in the fibroblast cell lines (CAFs and GFs) demonstrating their capacity to quickly respond to IL-1α. The epithelial cell lines were also responsive to IL-1α treatment, albeit to a lesser degree. (Figure 2E).

### Exogenous IL-1α does not promote secretion of cathepsin K

As we found that small amounts of cathepsin K in exosomes secreted by HSC-3 cells into media (Figure 2A), we assessed if IL-1α-driven cathepsin K overexpression could also increase extracellular cathepsin K. We measured cathepsin K levels in all of our cell lines, most of the cathepsin K remained intracellular, and in case of HSC-3, CAFs and GFs there was even a reduction in the amount of secreted cathepsin K to almost non-detectable levels after IL-1α treatment (Figure 2E).

### IL-1α has no effect on the invasiveness of HSC-3 cells in the myoma organotypic model

So far, it was established that cathepsin K levels, particularly in CAFs and GFs, could be affected by the presence of IL-1α. Therefore, we aimed to assess whether IL-1α-driven cathepsin K levels could affect HSC-3 invasiveness in the myoma organotypic invasion model. We designed six different experimental conditions, both including and excluding IL-1α supplementation and co-culture of OTSCC cells with GFs or CAFs. Our results confirm the HSC-3 cells intrinsic invasiveness, although there was no significant change in the invasion depth as a result of IL-1α treatment, or co-cultures with neither GFs nor CAFs (Figure 3).

**Figure 3.**
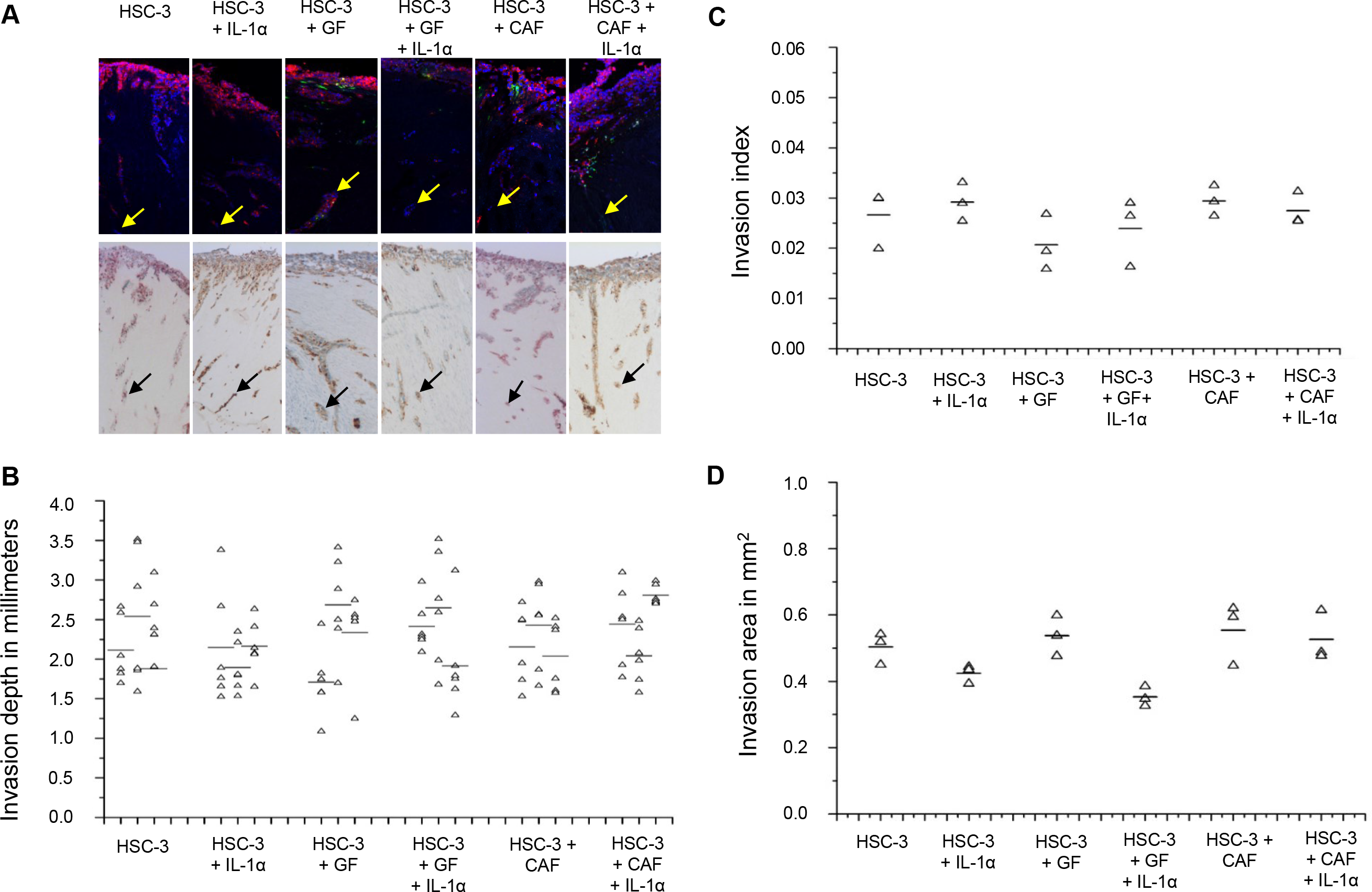
HSC-3 cells cultured in the myoma organotypic model present similar invasion depth and pattern, regardless of experimental conditions. **(A)** Arrows point to the deep invading OTSCCs in representative images of the myoma tissue, both in HSC-3 fluorescently labeled and in pancytokeratin immunostained cells. **(B)** The HSC-3 cells had similar invasion depth when cultured either together or separate from either GFs or CAFs. Also, in experimental conditions where IL-1α was added to culture media, there was no observed effect on HSC-3 invasiveness. The invasion depth shown was measured in fluorescent sample images and was shown to be consistent with the measurement in immunostained samples. Due to sample size, all samples were measured in six sections, each represented by a triangle. **(C)** The invasion index (calculated as 1-[non-invaded area/total area]) was found to be similar for all of the experimental conditions tested. Each triangle represents an average of all replicated of experimental groups, with a horizontal dash representing the average value. **(D)** The invasion area was similar for all samples analyzed. Each triangle represents an average of all replicated of experimental groups, with a horizontal dash representing the average value.

## DISCUSSION

The role of cathepsin K outside of bone extracellular remodeling is still relatively unknown. In spite of being virtually absent from normal tissue and present in both cancer cells and tumor stroma, this differential expression has, so far, shed only a little light on its function in the TME. With its powerful collagenase activity, it is possible to argue for an intense role of cathepsin K in TME matrix turnover, similar to wound healing and inflammatory processes (17, 18). However, cathepsin K presence in carcinoma cells points to a more complex dynamic, where the cross-talk between cancer and TME cells is of paramount importance to the regulation of its expression. Our efforts aimed to provide some insight into the relevance of IL-1α in the OTSCC microenvironment.

In our *in vivo* findings, we have extensively described the concentration of cathepsin K close to the cellular membrane (9). This membranous expression pattern, although not relevant for prognosis, suggested that OTSCC cells might have the ability to secrete cathepsin K-containing vesicles. Secretion of cathepsin K into the extracellular environment outside the bone tissue has already been reported in thyroid tissue (19, 20). Although the membranous localization is suggestive of cathepsin K secretion and the presence of cathepsin K in purified exosomes from HSC-3 cells, the immunostaining results did not show extracellular cathepsin K in the TME. In our *in vitro* experiments, neither inducing lysosomal purging nor stimulating cathepsin K synthesis has resulted in significant extracellular cathepsin K production. In PMA treated cells, cathepsin K containing vesicles seemed to scatter towards the cell membrane, similarly to what was seen in the *in vivo* samples. However, at any time point followed, there was no clear evidence of cathepsin K secretion. These findings are consistent with the ELISA experiments, where basal cathepsin K levels in cell culture media were nearly undetectable in IL-1α-treated OTSCCs cells while at the same time intracellular cathepsin K increased. This also points to the fact that cathepsin K is mostly not secreted by OTSCC cells, but small amounts may be released through exosomes and play a part in inter cellular signaling.

Our results established the differential levels of cathepsin K and IL-1α in epithelial and mesenchymal cells, with IL-1α levels higher in HSC-3, IHMK and HMK than in CAFs and GFs. IL-1α treatment had greater effect on the protein levels of cathepsin K in CAFs and GFs than in epithelial cell lines. These results confirm the findings by Xie *et al.* where IL-1α from skin cancer cells drove increased expression of cathepsin K in fibroblasts *in vitro* (5). It is also important to note that while CAFs and GFs were more responsive to IL-1α treatment, cathepsin K levels in the epithelial cells were also modestly increased. In our previous publication, we described how cathepsin K was present in OTSCC cells in tissue sections, and virtually absent in normal oral mucosal cells (9). Our *in vitro* results complement those findings, suggesting that cathepsin K may have been present in normal mucosa, although in small, undetected amounts. The same is true when healthy gingival tissue is compared with periodontitis-affected tissue. Normal tissue has very low numbers of cathepsin K positive cells, while the inflamed tissue presents increased cathepsin K immunoreactivity in oral fibroblasts (18).

Tumor cell invasion is an extremely important OTSCC hallmark, being essential for tumor progression. The invasion process involves an intricate collection of cellular processes, which will result in cancer cell detachment from the primary tumor, remodeling of the extracellular matrix and invasion of neighboring tissues and structures (21). As a powerful collagenase, cathepsin K’s putative role in OTSCCs is to aid both carcinoma and associated host cells in the invasion process, with both cancer cells themselves and stromal cells showing overexpression and significant remodeling of the TME (6,22,24–25).

Although, in our in vivo samples, there was evident overexpression of cathepsin K in both the OTSCC cells and the TME, with significance for prognosis, in our current investigation, in vitro, we found no differences in the invasion ability of HSC-3 cells neither as a result of IL-1α-derived cathepsin K overexpression nor after co-culture with oral fibroblasts.

We interpret the lack of increased invasion by HSC-3 cells, due to the fact that they are already exceedingly invasive when cultured in the myoma organotypic model, the results suggest that the increase in cathepsin K by IL-1α in both OTSCC and fibroblast cell lines is of little consequence to this process. We also postulate that cathepsin K activity is being modulated indirectly by other signaling pathways and by the degradation of myoma tissue: a highly complex, hypoxic and enriched environment (23) That assumption is supported by our previous findings, where it was shown that HSC-3 cells knocked-down for cathepsin K had a significant decrease in invasive potential, when compared to controls (9), demonstrating that cathepsin K was important for invasion.

In conclusion, our results show that IL-1α induces cathepsin K in epithelial and mesenchymal cell lines in vitro. Our findings also suggest that, in OTSCC cells, subcellular compartments, although not secreted, can quickly mobilize cathepsin K. The role for cathepsin K intracellular movement is still unknown; however, we can assume that it may be an important action by the cancer cells to aid in their survival and invasion throughout the tumor progression.

## ACKNOWLEDGEMENTS

This study was supported by grants from Oulu University KEVO, the Sigrid Juselius Foundation, the Finnish Dental Society Apollonia and FAPESP grant 13/05822-9. We thank Ms. Sanna Juntunen and Tanja Kuusisto for expert technical assistance. We also thank Dr. Veli-Pekka Ronkainen for assisting with immunofluorescence imaging.

